# *Fusobacterium sphaericum sp. nov.*, isolated from a human colon tumor, adheres to colonic epithelial cells and induces IL-8 secretion

**DOI:** 10.1101/2023.06.16.545380

**Authors:** Martha A. Zepeda-Rivera, Yannick Eisele, Alexander Baryiames, Hanrui Wu, Claudia Mengoni, Gianmarco Piccinno, Elsa F. McMahon, Kaitlyn D. LaCourse, Dakota S. Jones, Hans Hauner, Samuel S. Minot, Nicola Segata, Floyd E. Dewhirst, Christopher D. Johnston, Susan Bullman

## Abstract

Cancerous tissue is a largely unexplored microbial niche that provides a unique environment for the colonization and growth of specific bacterial communities, and with it, the opportunity to identify novel bacterial species. Here, we report distinct features of a novel *Fusobacterium* species, *F. sphaericum sp. nov.* (*Fs*), isolated from primary colon adenocarcinoma tissue. We acquire the complete closed genome and associated methylome of this organism and phylogenetically confirm its classification into the *Fusobacterium* genus, with *F. perfoetens* as its closest neighbor. *Fs* is phenotypically and genetically distinct, with morphological analysis revealing its coccoid shape, that while similar to *F. perfoetens* is rare for most *Fusobacterium* members. *Fs* displays a metabolic profile and antibiotic resistance repertoire consistent with other *Fusobacterium* species. *In vitro, Fs* has adherent and immunomodulatory capabilities, as it intimately associates with human colon cancer epithelial cells and promotes IL-8 secretion. Analysis of the prevalence and abundance of *Fs* in >20,000 human metagenomic samples shows that it is a low-prevalence member within human stool with variable relative abundance, found in both healthy controls and patients with colorectal cancer (CRC). Our study sheds light on a novel bacterial species isolated directly from the human CRC tumor niche, and given its interaction with cancer epithelial cells suggests that its role in human health and disease warrants further investigation.

## Introduction

In patients with colorectal cancer (CRC), unbiased genomic analyses have revealed an enrichment of *Fusobacterium* in CRC tumors relative to noncancerous colorectal tissue^1–4^. Previous work from our group and others demonstrated that within patient CRC tumors, *Fusobacterium* colonizes distinct regions with immune and epithelial functions supportive of cancer progression^5^, that it persists with the primary tumor during metastasis^6^, and that antimicrobial agents targeting these and other anaerobic bacterial species decrease cancer cell proliferation and tumor growth^6,7^. Furthermore, high *Fusobacterium* loads are associated with high-grade dysplasia^1^, advanced disease stage^8^, and poorer patient prognosis^9^. As tissue-associated microbiomes are not readily amenable to metagenomic analysis, due to the overabundance of host nucleic acids, characterization of tumor-infiltrating microbial communities is often restricted to amplicon-based sequencing approaches with limited phylogenetic resolution. To overcome this challenge, microbiological culture-based approaches with patient tissue specimens have reemerged as valuable tools to isolate members of tissue-associated microbiomes.

Through a culturing-based approach, we isolated, characterized, and sequenced the genome of a novel *Fusobacterium* species from the resected tumor of a treatment naïve female patient with microsatellite stable (MSS), stage III, right-sided colon cancer. Phylogenetic appraisal of this organism places this bacterium within the *Fusobacterium* genus but demonstrates that it is distinct from currently known species. Rather than the typical spindle-like, fusiforme morphologies associated with most *Fusobacterium* species, phenotypic analysis of this novel organism revealed a coccobacilloid shape. Acknowledging this distinct spherical cell-shape, we propose this species as *Fusobacterium sphaericum sp. nov.* (*Fs*). In this present study, we find that while the chromosomal structure and gene content of *Fs* is predominantly unique to this species, its metabolic profile and antibiotic resistance repertoire are consistent with other *Fusobacterium* members. Furthermore, our analyses indicate that its predicted virulence factors are related to metabolism, cell-cell adherence, and immunomodulation. Further, *in vitro* co-culture with human colon cancer epithelial cells confirmed *Fs* adherence and indicated that its presence significantly increases IL-8 secretion. Assessment of the prevalence and abundance of *Fs* in 21,212 human metagenomic samples shows that it is a low-prevalence member in human stool with variable relative abundance.

## Results

### Phylogenetic and morphologic characterization of a novel Fusobacterium species

*Fusobacterium* isolation was performed on a human colorectal cancer (CRC) tumor from a treatment naïve female patient with microsatellite stable (MSS), stage III, right-sided colon cancer. Using long-read <u>s</u>ingle-<u>m</u>olecule real-time (SMRT)^10^ sequencing we generated a complete and closed genome and methylome for isolate SB021, identifying two putative chromosomes (1.69 Mb and 0.57 Mb) and two additional extrachromosomal elements (1.59 Kb and 0.32 Kb) (**Fig. 1a, Supplementary Table 1, Supplementary Table 2**). BLASTn analysis of the 16S rRNA gene of SB021 confirmed its classification into the *Fusobacterium* genus with a top hit against *Fusobacterium perfoetens*, yet all related hits fell below the established 16S rRNA gene species similarity threshold of 98.65%^11^, suggesting that SB021 represents a novel *Fusobacterium* species (**Supplementary Table 3**). Comparative phylogenetics of SB021 and other *Fusobacterium spp.* genomes (**Table 1**), both by the 16S rRNA gene and a reference-free whole-genome approach^12^, further showed that SB021 forms a distinct branch within the same clade as *F. perfoetens* (**Fig. 1b-c**). To quantify the relatedness between SB021 and the publicly available *F. perfoetens* ATCC 29250 genome, we calculated the average nucleotide identity (ANI) and found it to be 75.27%, well below the established species threshold of 95-96%^11^, supporting the classification of SB021 as a novel *Fusobacterium* species. (**Fig. 1d, Supplementary Fig. 1, Supplementary Table 4**). GTDB-tk^13^ analysis further supported these observations, as it classified SB021 within the same clade as *F. perfoetens*, as part of “Fusobacterium_B_sp900541465” (**Supplementary Table 5**). This group is predominantly represented by metagenomically assembled genomes (MAGs) from uncultured samples, with a single uncharacterized isolate from a large-scale cultivation effort of the fecal microbiota in healthy Chinese individuals^14^.

**Fig. 1:**
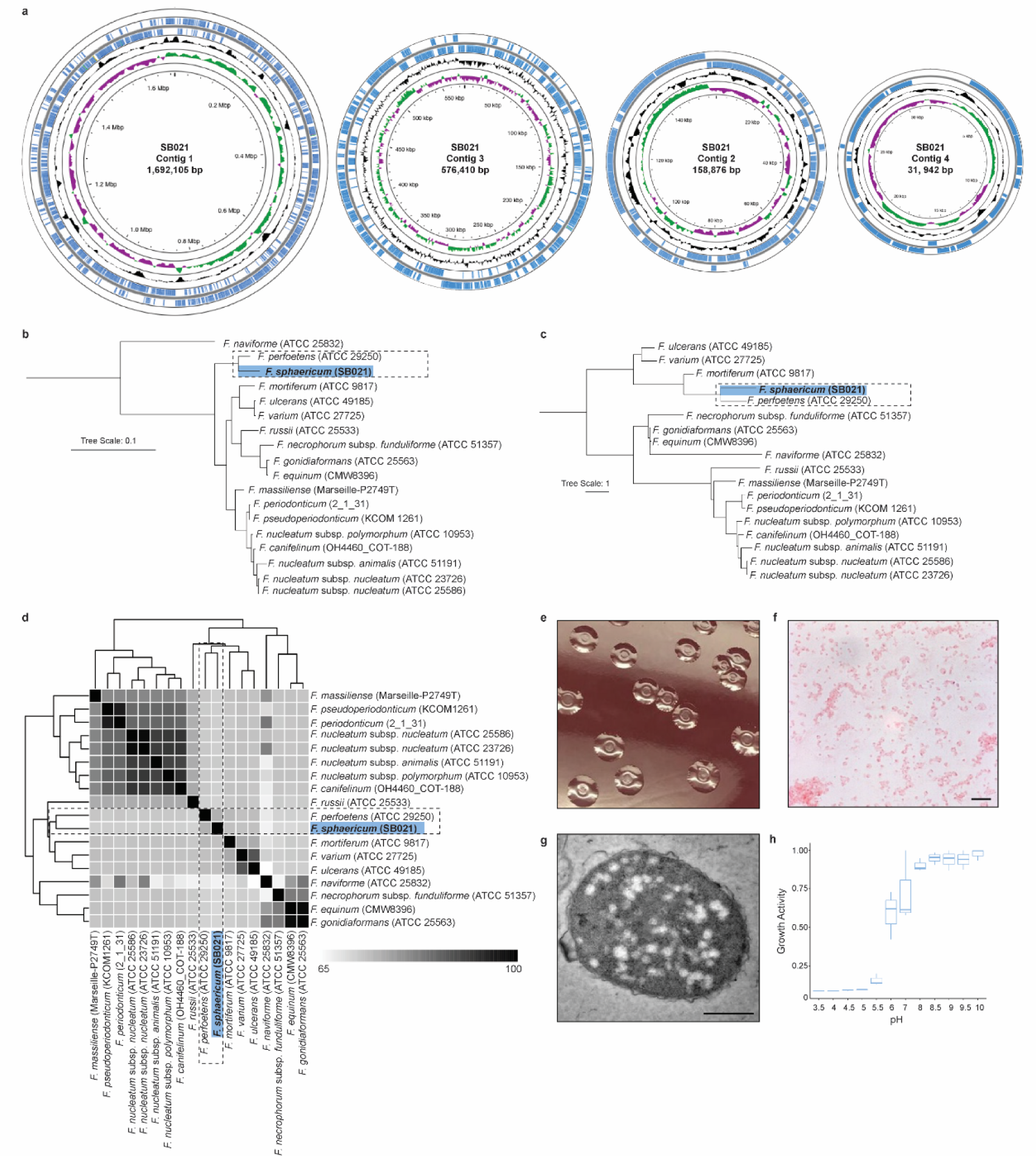
Taxonomic and Morphologic Analysis of *Fusobacterium sphaericum sp. nov.* (SB021) **a)** Proksee circular genome architecture of *F. sphaericum sp. nov.* (*Fs*) SB021 contigs. From outwards to inwards, the first and second circles show annotated coding sequences on positive strand and negative strand, respectively. Third circle indicates average GC content, with areas higher than average GC content as outward projections and areas lower than average as inward projections. Fourth circle shows GC skew, with above zero values in green and below zero values in purple. **b)** A 16S rRNA gene-based dendrogram illustrating the phylogenetic relationship between *Fs* (blue) and other *Fusobacterium spp*. **c)** kSNP^12^ maximum-likelihood whole-genome phylogenetic tree with *Fs* highlighted in blue. **d)** Clustered average nucleotide identity (ANI) matrix with *Fs* highlighted in blue. ANI values are reported in **Supplementary Table 4**. White to black scale indicates and ANI from 65% to 100%, respectively. **e)** Digital photography of *Fs* SB021 colonies grown on fastidious anaerobe agar enriched with 10% defibrinated horse blood. **f)** Bright-field microscopy of Gram-stained *Fs* SB021. Images acquired at 100X magnification with oil immersion lens. Image scale bar is 10 µm. **g)** Transmission electron micrographs of an *Fs* SB021 cell. Image scale bar is 500 µm. **h)** Plot indicates growth activity as measured in Biolog PM10 plates for *Fs* SB021. Data is normalized to the maximum value per replicate (n=3).

**Table 1:** Genomes used in this study. Table indicates each *Fusobacterium* genome used for comparative analyses in this study with their corresponding genome accession numbers, genome length, GC content, and number of contigs in the assembly noted.

The cellular morphology and pH tolerance of *Fusobacterium* cells is known to vary^15–17^. As SB021 is phylogenetically closely related to *F. perfoetens*, we assessed whether like *F. perfoetens*, SB021 cells have a short, ovoid coccobacillus morphology. Macroscopically, SB021 cells form distinct, vaguely round, raised opaque colonies with central indentations following 48 hours of anaerobic culture. The colonies are viscous and malodorous and measure 3-5 millimeters in diameter (**Fig. 1e**). Microscopically, SB021 cells are short, gram-negative coccobacilli, with an average length of 1.60μm and an average width of 1.25μm (**Fig. 1f**). Transmission electron microscopy further revealed a spherical shape and a granulated cytoplasm (**Fig. 1g**). These results suggest that members within this *Fusobacterium* clade share similar cellular morphology. Assessment for preferential growth pH indicated that SB021 is sensitive to pH below 5 and demonstrates a growth preference between pH 8 and pH 10 (**Fig. 1h**). Given the coccobacilloid morphology of SB021 cells, we propose this novel species, part of “Fusobacterium_B_sp900541465” as *Fusobacterium sphaericum sp. nov.* (*Fs*).

### *Fs* genomic attributes predict metabolic and virulence capabilities

As significant genomic heterogeneity has been reported within the *Fusobacterium* genus, with genome size, architecture, and content differing between species of varying pathogenic capabilities^17,18^, we sought to characterize the genomic content of *Fs.* To compare the genetic content of *Fs* to other *Fusobacterium spp*., we implemented the <u>a</u>nalysis and <u>v</u>isualization platform for <u>‘o</u>mics data (Anvi’o) workflow for microbial pangenomics^19^. This genus-level pangenomic analysis allows us to identify all genes present in the *Fusobacterium* genus (“pangenome”) and discern between gene content that is shared amongst ≥95% of species (“core genome”), is shared amongst subsets of species (“accessory genomes”) or is unique to individual members (“singletons”). Across eighteen representative *Fusobacterium* genomes (**Table 1**), only 4.9% of gene clusters compose the core (576/11,710), indicative of the extreme genetic heterogeneity across this genus (**Fig. 2a**). *Fs* harbored 18.7% of identified gene clusters (2,192/11,710), but of these, 31.8% (697/2,192) were unique to *Fs*, while only 6.8% (150/2,192) were shared with its closest phylogenetic neighbor, *F. perfoetens* (**Fig. 2b**).

**Fig. 2:**
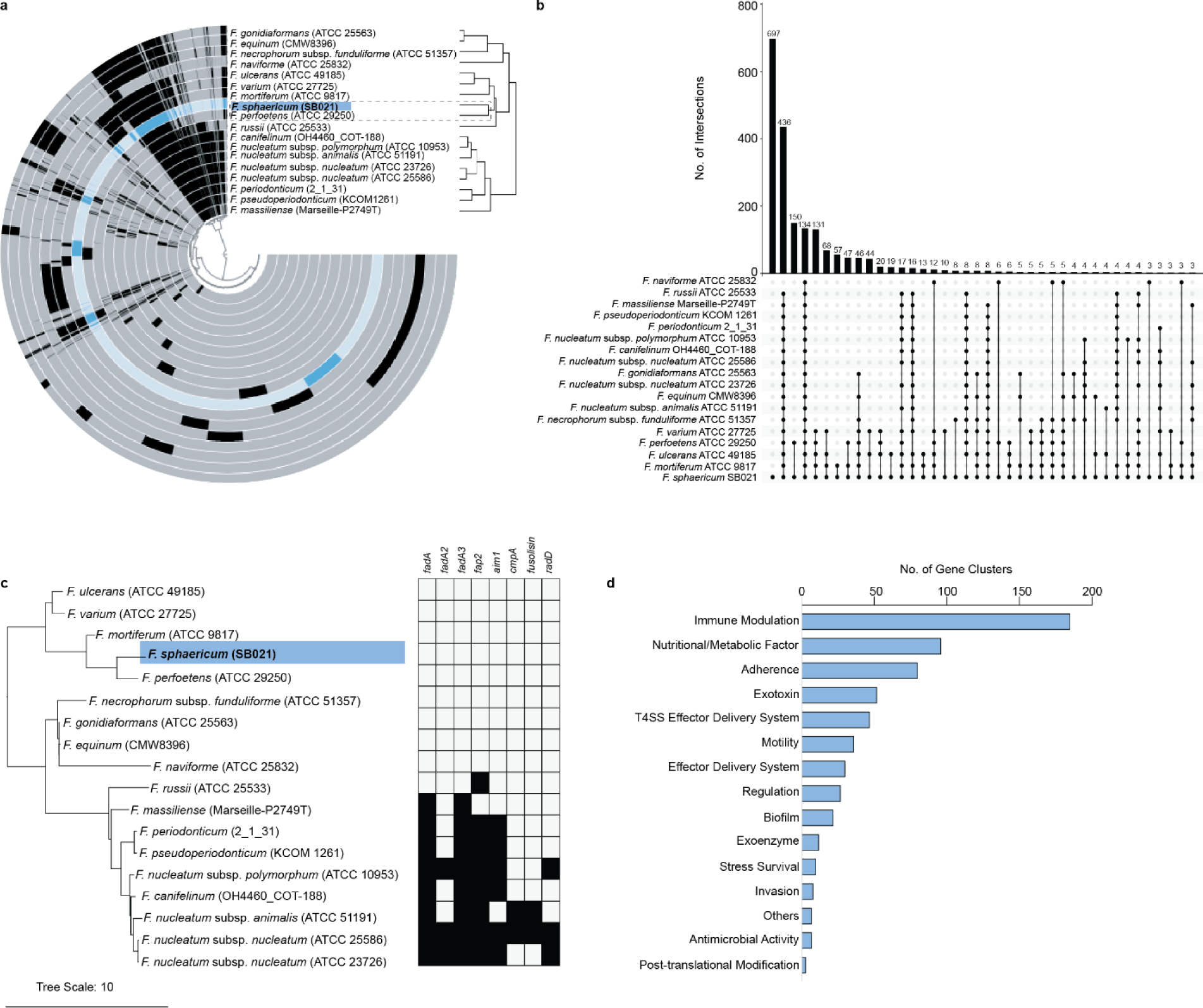
*Fusobacterium sphaericum sp. nov.* (SB021) Gene Content Analysis. **a)** Anvi’o pangenomic analysis^19^ of *F. sphaericum sp. nov.* (*Fs*) SB021 (blue) as compared to seventeen other *Fusobacterium spp.* genomes (**Table 1**). Each circle represents an individual genome, with identified gene clusters in blue (SB021) or black (other *Fusobacterium spp*.). Genomes are ordered by ANI clustering (Fig. 1d). **b)** Upset plot depicting the presence of gene clusters identified in *Fs* across other *Fusobacterium spp.* genomes. Bar height (top) indicates number of gene clusters shared between combinations of genomes (bottom). **c)** Presence (black) versus absence (white) plot of canonical virulence factors across *Fusobacterium spp.* genomes, ordered by a kSNP^12^ maximum-likelihood whole-genome phylogenetic tree with *Fs* highlighted in blue. **d)** *Fs* anvi’o-identified gene clusters were analyzed using the virulence factor database (VFDB)^33^. Graph shows the number of gene clusters in each of the top functional categories of putative virulence factors (**Supplementary Table 5**).

As the genetic content of *Fs* is predominantly unique to this species, we assessed the 2,192 identified gene clusters. KEGG ortholog analysis^20^ revealed that mapped gene clusters (50.8%) were predominantly involved in metabolic functions and pathways (**Supplementary Table 6**). Primary metabolic pathways assessment via gutSMASH^21^ indicated shared predicted metabolic capabilities between *Fs* and other *Fusobacterium* genomes (**Supplementary Fig. 2**). *In vitro* metabolic profiling via the API 20A Gallery System showed that *Fs* ferments the monosaccharides glucose and mannose, the disaccharide lactose, the trisaccharide raffinose, and the coumarin glucoside esculin (**Supplementary Fig. 3a**). Glucose, mannose, and raffinose metabolism were also observed via the Biolog Anaerobe Identification Test Panel (AN Plate) (**Supplementary Fig. 4**). Enzymatic profiling via the API ZYM panel additionally showed positive activity for alkaline phosphatase, C4-esterase, C8-esterase-lipase (weak), acid phosphatase (weak), napthol-AS-BI-phosphohydrolase (weak) and α-Galactosidase (strong) (**Supplementary Fig. 3b**). This strong α-Galactosidase activity may be a distinct feature of *Fs*, as activity for this enzyme has only been observed weakly in *F. mortiferum* and *F. necrogenes*^22^.

Notably, *Fs* harbors nineteen open reading frames encoding putative multidrug resistance proteins (**Supplementary Table 1**) and further analysis with the comprehensive antibiotic resistance database (CARD)^23^ identified 183 putative antibiotic resistance genes (**Supplementary Table 7**), suggesting that *Fs* is resistant to a suite of antibiotics. Our group has previously demonstrated that antibiotic-mediated microbiota modulation, targeting *Fusobacterium* and other anaerobes, decreases cancer cell proliferation and tumor growth^6^. As such, we sought to characterize the antibiotic response of *Fs. In vitro* antibiotic susceptibility testing demonstrated that *Fs* is susceptible to various classes of antibiotics, including metronidazole (MIC 0.502 μg/ml; nitroimizadole class), colistin (MIC 0.875 μg/ml; polymyxin class), and penicillin (MIC 0.079 μg/ml; beta-lactam class). Consistent with other *Fusobacterium spp*., *Fs* is resistant to erythromycin (macrolide class) and vancomycin (glycopeptide class) antibiotics. Curiously, *Fs* susceptibility to aminoglycosides varied, with susceptibility to gentamycin (MIC 62.2 μg/ml) and resistance to streptomycin and kanamycin (**Table 2**).

**Table 2:** *F. sphaericum sp. nov.* antibiotic resistance profile. Table shows results for *F. sphaericum sp. nov.* susceptibility or resistance to nine tested antibiotic compounds.

Given that *Fs* was isolated directly from a patient CRC tumor, we next queried the presence of virulence factors important for host-bacterial interactions. We found that *Fs* lacks known *Fusobacterium* virulence factors^24–32^ (**Fig. 2c**) but of the 2,192 identified gene clusters, 28.4% of them are homologous to known virulence factors in other bacterial pathogens^33^ (**Supplementary Table 6**). Of this subset, the majority function to modulate the immune system (29.7%), to scavenge and metabolize nutritional sources (15.4%), or to adhere to host cells (12.9%) (**Fig. 2d**). This suggests that *Fs* has genetic attributes consistent with facilitating eukaryotic cell interactions and modulation of immune responses.

### *Fs* adheres to colon cancer epithelial cells and increases IL-8 secretion

To evaluate the interaction between *Fs* and colon epithelial cells, we co-cultured *Fs* SB021 with cell lines derived from human colorectal adenocarcinomas, HCT116 and HT-29. As *Fusobacterium nucleatum* subsp. *animalis* (*Fna*) has previously been reported to adhere to and invade cancer epithelial cells^6^, we used *Fna* SB010 as a positive control. Confocal laser scanning microscopy of fluorescent immunocytochemically stained cells demonstrated that SB021 strongly adheres to epithelial cell surfaces in chain-like clusters with both HCT116 and HT-29 (**Fig. 3a**). However, we did not observe intracellular SB021, which suggests that, unlike SB010, SB021 adheres to but does not invade colonic epithelial cells.

**Fig. 3:**
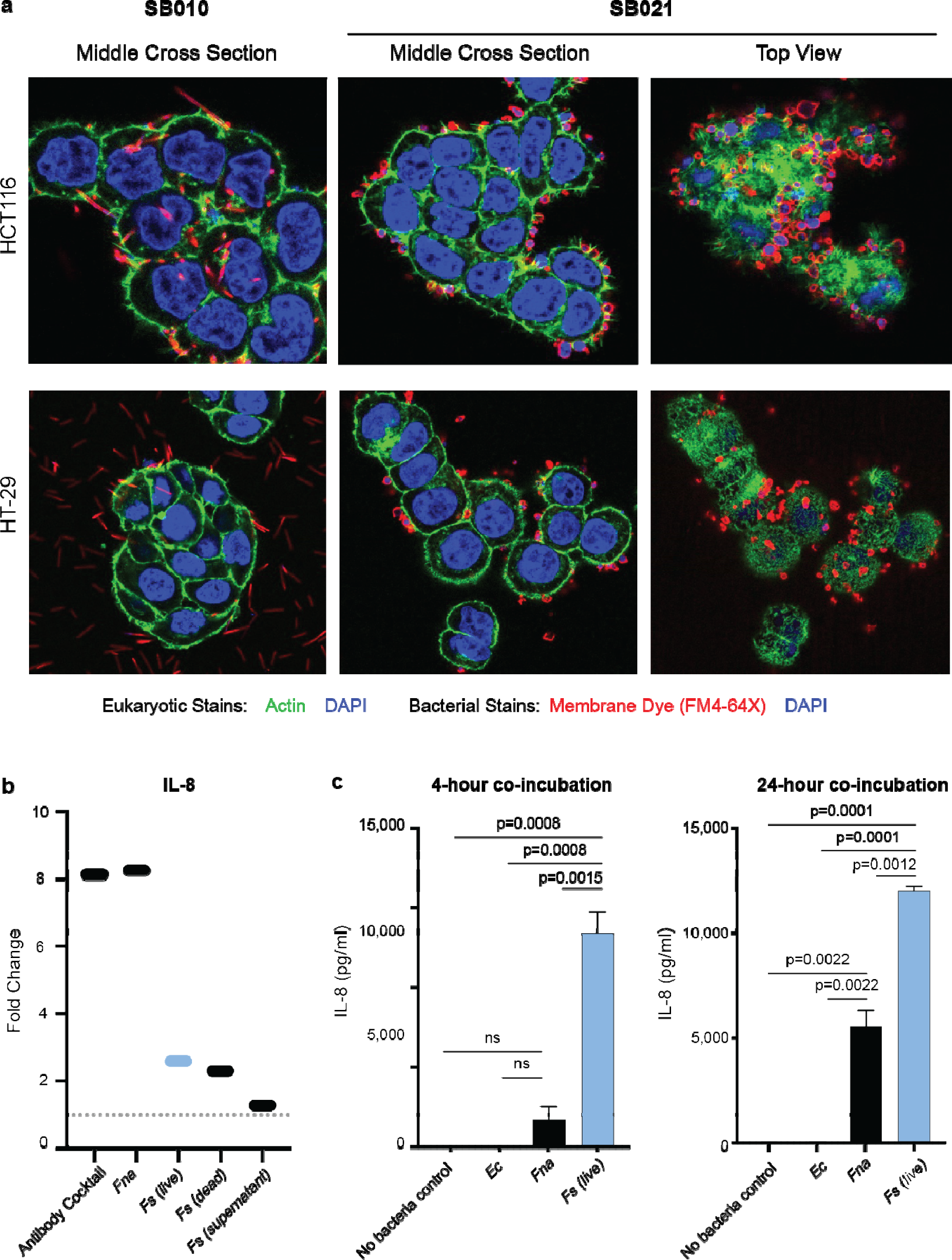
*Fusobacterium sphaericum sp. nov.* adheres to colonic epithelial cells and stimulates IL-8 secretion. Confocal laser scanning microscopy of colon cancer epithelial cell lines, HCT116 and HT-29, co-incubated with *Fusobacterium nucleatum* subsp. *animalis* (*Fna*) SB010 or *F. sphaericum sp. nov.* (*Fs*) SB021. Bacterial cell staining: FM4-64X (red) and DAPI (blue). Eukaryotic cell staining: DAPI (blue), actin (green). **b)** Comparison of IL-8 secretion measured via the Multi-Analyte ELISArray from HCT116 cells co-incubated with different *Fusobacterium* species or antibody cocktail control, normalized to a no-bacteria control. Bacterial strains used included SB010 and SB021 live cells or oxygen-exposed dead cells. An additional co-incubation with SB021 supernatant alone was also performed. Additional cytokine and chemokine measurements are shown in **Supplementary Fig. 5**. **c)** Quantitative levels of IL-8 secretion measured via the Single-Analyte ELISArray from HCT116 cells incubated alone or co-incubated with *Escherichia coli* TOP10, SB010, or SB021 at 4-hour and 24-hour time points. Statistical analysis performed via a one-way ANOVA. For **a-c**, all co-incubations were at a multiplicity of infection (MOI) of 100.

To investigate whether this physical interaction between *Fs* and colonic epithelial cells modulates eukaryotic immune responses, we assessed chemokine and cytokine induction in HCT116 cells post co-culturing. In the presence of SB021 cells, we observed an increase in various pro-inflammatory chemokines, most notably IL-8, occurring with both live (2.59-fold increase) and dead (2.29-fold increase) bacterial cells (**Fig. 3b, Supplementary Fig. 5**), indicating that these immunomodulatory capabilities of SB021 are independent of viability. IL-8 expression is elevated in colonic tissue of patients with CRC^34^, enhances colon cancer cell growth and migration to the liver and lungs^35,36^, and has previously been reported to be stimulated by the presence of other Fusobacterium species including *Fna* and *F. nucleatum subsp. nucleatum* (*Fnn*)^37^. Prior genetic analysis in *Fnn* indicated that the stimulation of IL-8 expression is dependent on Fap2-mediated cell adhesion^37^. To assess whether physical interaction between *Fs* and colonic epithelial cells is required for IL-8 secretion, we co-incubated SB021 cell-free supernatant with HCT116 cells and detected a 1.27-fold increase in IL-8 (**Fig. 3b**). These results suggest that unlike other Fusobacterium species, direct bacterial-eukaryotic cell interactions with *Fs* are not required to stimulate secretion of IL-8 and other pro-inflammatory chemokines and cytokines (**Fig. 3b, Supplementary Fig. 5**). Furthermore, quantitative comparison of IL-8 production in the presence of SB021 or SB010 further showed that co-incubation with SB021 stimulated higher IL-8 levels at both 4-hour (SB021: average 13,403 pg/ml; SB010: average 1,732 pg/ml) and 24-hour (SB021: average 12,004 pg/ml; SB010: average 5,551 pg/ml) time points (**Fig. 3c**).

### *Fs* is a low prevalence member in human stool

Given that *Fs* is a novel *Fusobacterium* species with adherent and immunomodulatory capabilities, we next sought to determine its prevalence in human specimens from various body sites. The presence of *Fs* SB021 and its nearest phylogenetic neighbor, *F. perfoetens* ATCC 29250, was assessed across 21,212 publicly available human metagenomic samples from the human oral cavity, skin, vagina, airways, breastmilk, nasal passage, and stool (**Fig. 4a, Supplementary Table 8**). Results indicated that SB021 could be detected in stool samples at a low prevalence (147/21,382=0.687%), with a range of relative abundances from 0.0005%-17.829% when detected (**Fig. 4b**). Of the 147 samples in which SB021 could be detected, 31 originated from cohorts comparing patients with CRC to healthy controls. Nevertheless, no significant difference in SB021 prevalence (p=0.1936, two sample Z-test) (**Fig. 4c**) or abundance (p=0.8124, Welch’s T-test) (**Fig. 4d**) was observed based on disease status in those cohorts. In contrast, ATCC 29250 was detected in only a single stool sample with a low relative abundance of 0.00468% (**Supplementary Table 8**). Collectively, these findings shed light on a novel bacterial species isolated from the CRC tumor niche, which is also detectable within the human intestinal microbiota.

**Fig. 4:**
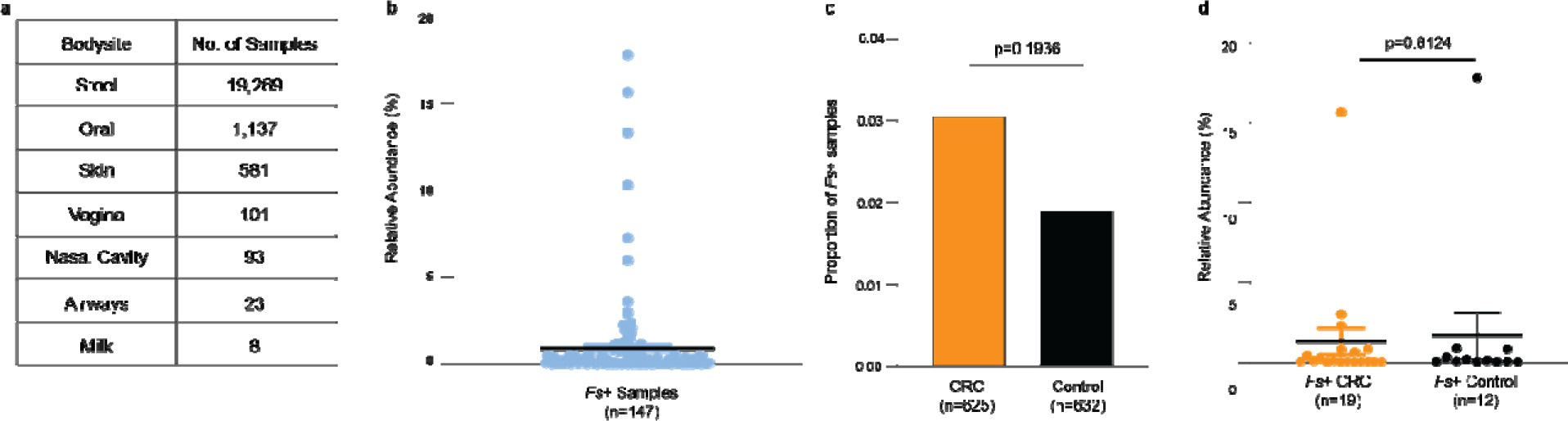
*Fusobacterium sphaericum sp. nov.* is present in human stool. The relative abundance of *F. sphaericum sp. nov.* (*Fs*) SB021 was assessed in **a)** 21,212 metagenomic samples from a various human body sites. **b)** Graphs show *Fs* percent relative abundance in samples detected. Black line indicates mean. **c-d)** In datasets comparing patients with CRC to healthy controls **c)** graph shows proportion of samples in which *Fs* was detected. Statistical analysis performed via two-sample Z test. **d)** Graph shows *Fs* percent relative abundance in samples detected for patients with CRC or healthy controls. Black line indicates mean. Statistical analysis performed via Welch’s T-test.

## Discussion

Owing to the abundance of host-associated nucleic acids in complex specimens, tissue-associated microbiome analysis is currently restricted to amplicon-based sequencing. As such, applying traditional culturing-based approaches to tumor specimens provides a valuable opportunity to expand our understanding and classification of tumor-infiltrating microbes^38^. Here, we report the isolation and characterization of a novel *Fusobacterium* species, *F. sphaericum sp. nov*. (*Fs*), from a primary colon adenocarcinoma of a female patient. Phylogenetic and pangenomic analyses reveal that *Fs* is most closely related to *F. perfoetens* (ATCC 29250), but support that *Fs* is a distinct species with predominantly unique gene content (**Fig. 1**, **Fig.2**).

Motivated by the putative cell-cell adherent and immunomodulatory capabilities of *Fs* genetic content (**Fig. 2d**), we investigated the interactions between *Fs* and colonic epithelial cells *in vitro*. Transmission electron microscopy and confocal microscopy of fluorescent immunocytochemically stained cells from *Fs* co-culture with human colon epithelial cell lines demonstrated that *Fs* adheres to eukaryotic epithelial cells but does not appear to have invasive capabilities, in contrast to other *Fusobacterium* species, such as *F. nucleatum* subsp. *animalis* (*Fna*) (**Fig. 3a**). *Fs* cells furthermore induced eukaryotic IL-8 secretion from epithelial cells at higher levels than *Fna* (**Fig. 3b-c**). Recent studies have demonstrated increased secretion of IL-8 in colorectal cancer (CRC) tissue and liver metastases^34,35^ with a significant upregulation in the presence of other *Fusobacterium* species including *Fna* and *F. nucleatum* subsp. *nucleatum (Fnn)*^37^. However, in *Fnn* induction of IL-8 secretion is dependent on Fap2-mediated adhesion to host cells^37^. As *Fs* lacks Fap2 (**Fig. 2c**) and was not observed to invade colonic epithelial cells (**Fig. 3a**), further studies to identify the *Fs* factors that induce IL-8 secretion are warranted. As *Fs* induction of IL-8 secretion was comparable independent of bacterial cell viability, and to a lower extent with cell-free supernatants (**Fig. 3b**), it is possible such factors are secreted or present in outer membrane vesicles. Demonstrating that *Fs* induces IL-8 secretion adds a novel species to a growing body of evidence for a distinct role of microbes contributing to a pro-inflammatory environment within human tissues, including tumors.

We demonstrate that in human metagenomic samples (n=21,212) *Fs* is also a member of the human stool microbiota in both healthy individuals and patients with CRC (**Fig. 4**). Although no significant difference in prevalence or abundance based on health status was observed, this could be due to the small number of *Fs*^+^ samples (n=31). Future genomic studies including *Fs* will be required to reveal its relationship with other microbial members and decipher its impact on the human host. Whether the presence of *Fs* in colorectal patients may contribute to a model of dysbiosis and chronic bacterial infection in the context of CRC requires further investigation in animal models and within longitudinal studies of large human cohorts.

The closest phylogenetic neighbor of *Fs* SB021, *F. perfoetens* ATCC 29250, was only detected in a single human stool sample. It is noteworthy that the first described isolate of *F. perfoetens*, acquired in 1900 from the diarrhea of infants, was unfortunately lost to history. As a result, subsequent isolates from a horse cecum (strain CC1) and piglet feces (strain ATCC 29250) were classified as the same species as the original 1900 isolate, but based only on cellular morphology and metabolism, as sequencing technologies for phylogenetic classifications were unavailable at the time^16^. Although this species underwent various name changes, ATCC 29250 was designated as the type strain for species *F. perfoetens* and remains the only available isolate in existence^16^. The very low detection of *F. perfoetens* ATCC 29250 across human metagenomic samples is likely reflective of its swine origin and perhaps suggests that the original 1900 human stool isolate was a phylogenetically distinct microbe despite morphological similarities. With its similar coccobacilloid morphology, it is possible that *Fs* SB021, isolated from a human colonic tumor, is a phylogenetic representative of the original 1900 human isolate, distinct from *F. perfoetens* ATCC 29250.

In conclusion, in this study we have emphasized and demonstrated the importance of reductionist approaches, such as traditional microbial culturing, to isolate and identify members of tissue-associated microbiota which are not yet amenable to metagenomics. Next generation sequencing techniques, such as long-read SMRT-Sequencing^10^, allows for the assembly of high-quality, closed, and complete genomes from which accurate phylogenetic characterization, putative functions and virulent capabilities can be predicted through comparative genomic analyses. Deciphering the role of identified novel microbial species within tumor-associated bacterial communities on disease outcomes remains crucial for informing future preventative and therapeutic interventions.

## Supplementary Figures

**Supplementary Fig. 1:**
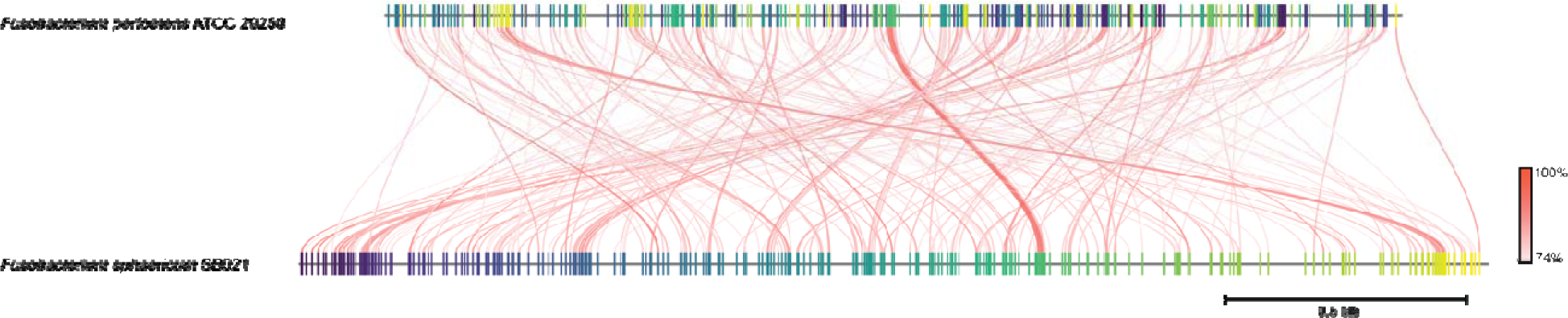
Genomic architecture of *Fusobacterium sphaericum sp. nov.* SB021 as compared to *Fusobacterium perfoetens* ATCC 29250. Visualization of fastANI alignment between *Fusobacterium sphaericum sp. nov.* SB021 and its nearest phylogenetic neighbor, *Fusobacterium perfoetens* ATCC 29250, where each red line segment denotes a reciprocal mapping of the 275 orthologous matches between the two genomes, indicating evolutionarily conserved regions.

**Supplementary Fig. 2:**
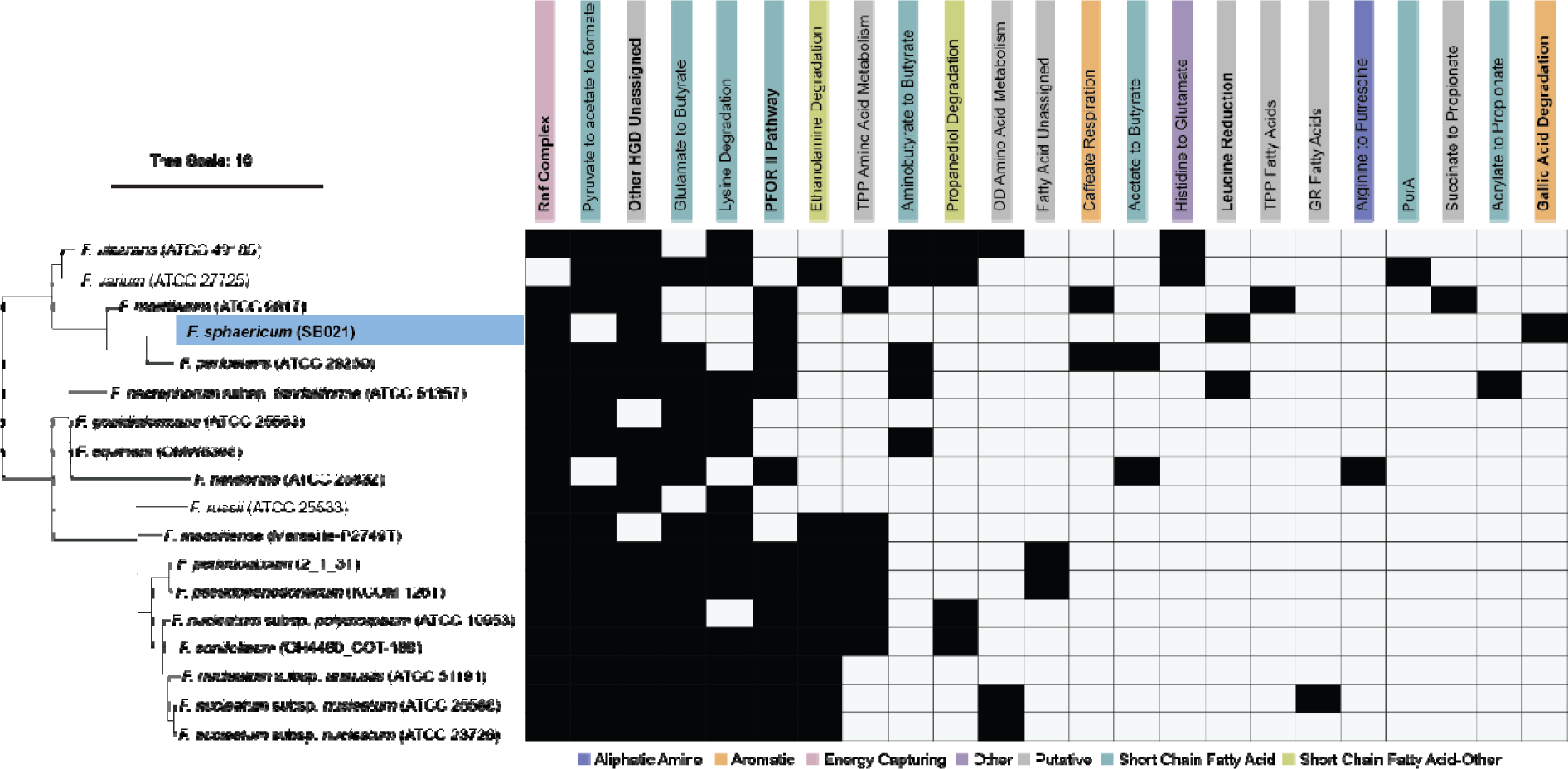
Primary metabolic pathways in *Fusobacterium sphaericum* SB021. Presence (black) versus absence (white) plot of primary metabolic pathways identified by gutSMASH^21^ across *Fusobacterium spp.* genomes, ordered by a kSNP^12^ maximum-likelihood whole-genome phylogenetic tree with *F. sphaericum sp. nov.* SB021 highlighted in blue.

**Supplementary Fig. 3:**
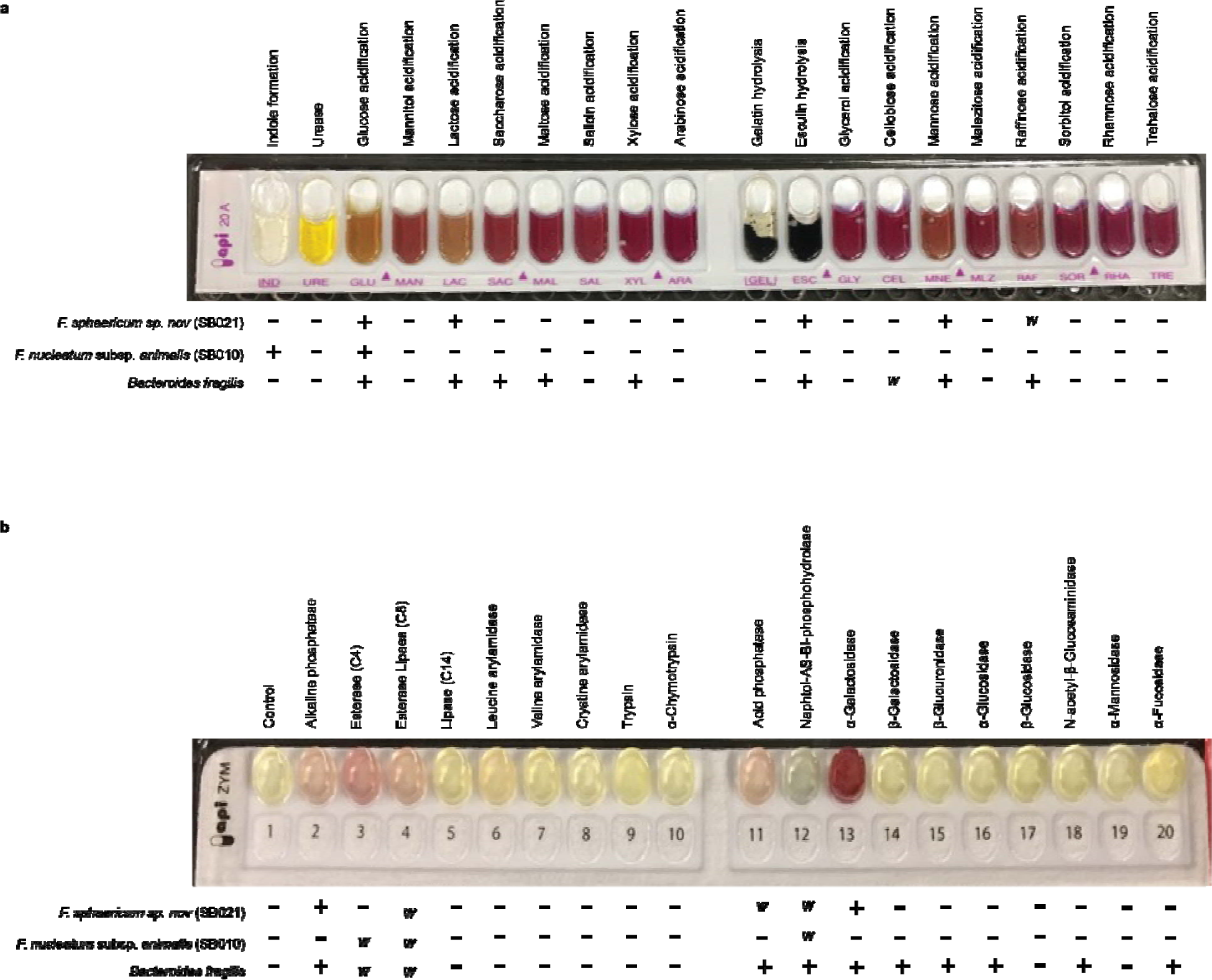
API metabolic characterization of *Fusobacterium sphaericum sp. nov*. *F. sphaericum sp. nov.* (*Fs*) **a)** metabolic and **b)** chemotaxonomic characteristics assayed by API 20A and API ZYM detection kits, respectively. Images are representative results for *Fs* SB021. Results for *Fs* SB021, *Fusobacterium nucleatum* subsp. *animalis* SB010, and *Bacteroides fragilis* are summarized below each API strip, where (+) indicates a positive result, (-) a negative result, and “*w*” a weakly positive result.

**Supplementary Fig. 4:**
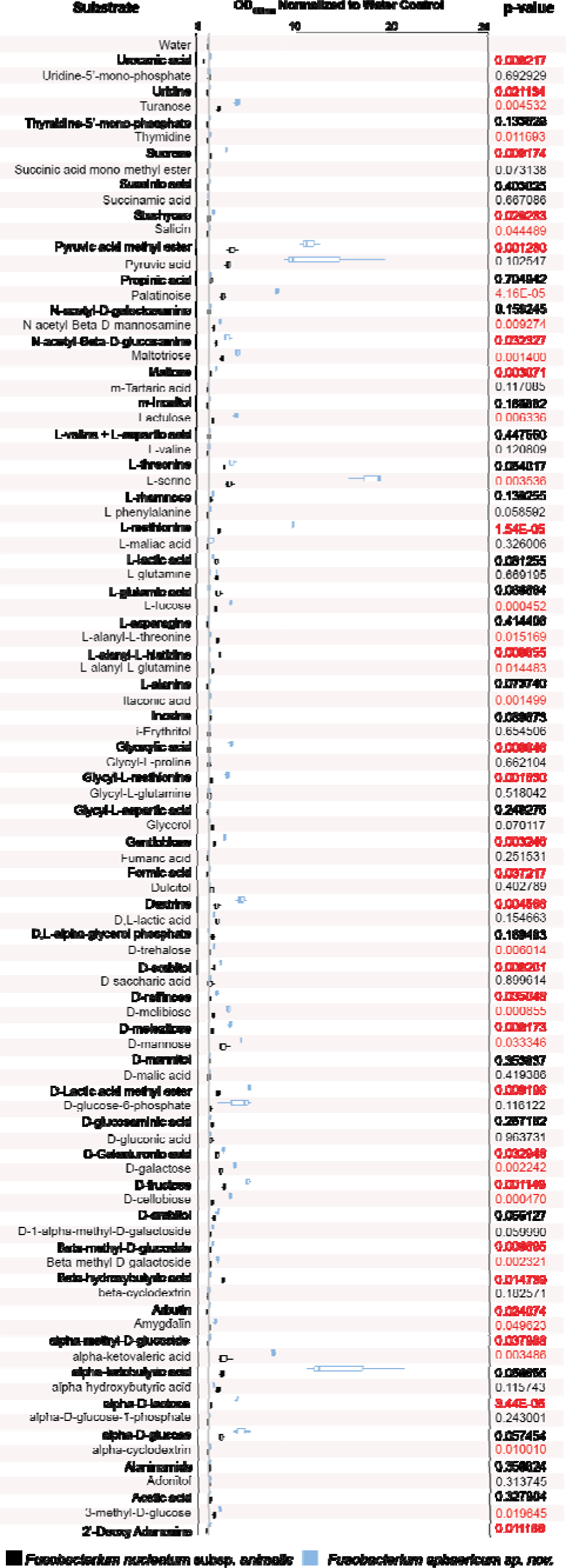
Biolog anaerobe identification test panel characterization of *F. sphaericum sp. nov.* substrate utilization. *F. sphaericum sp. nov.* substrate utilization assayed by Biolog Anaerobe Identification Test Panel (AN) assay plate. Graphs show results for *F. sphaericum sp. nov.* SB021 (blue) and *Fusobacterium nucleatum* subsp. *animalis* (*Fna*) SB010 (black) normalized to internal water control, in triplicate. Statistical analysis performed via a two-sided T-test. All p-values less than 0.05 are highlighted in red.

**Supplementary Fig. 5:**
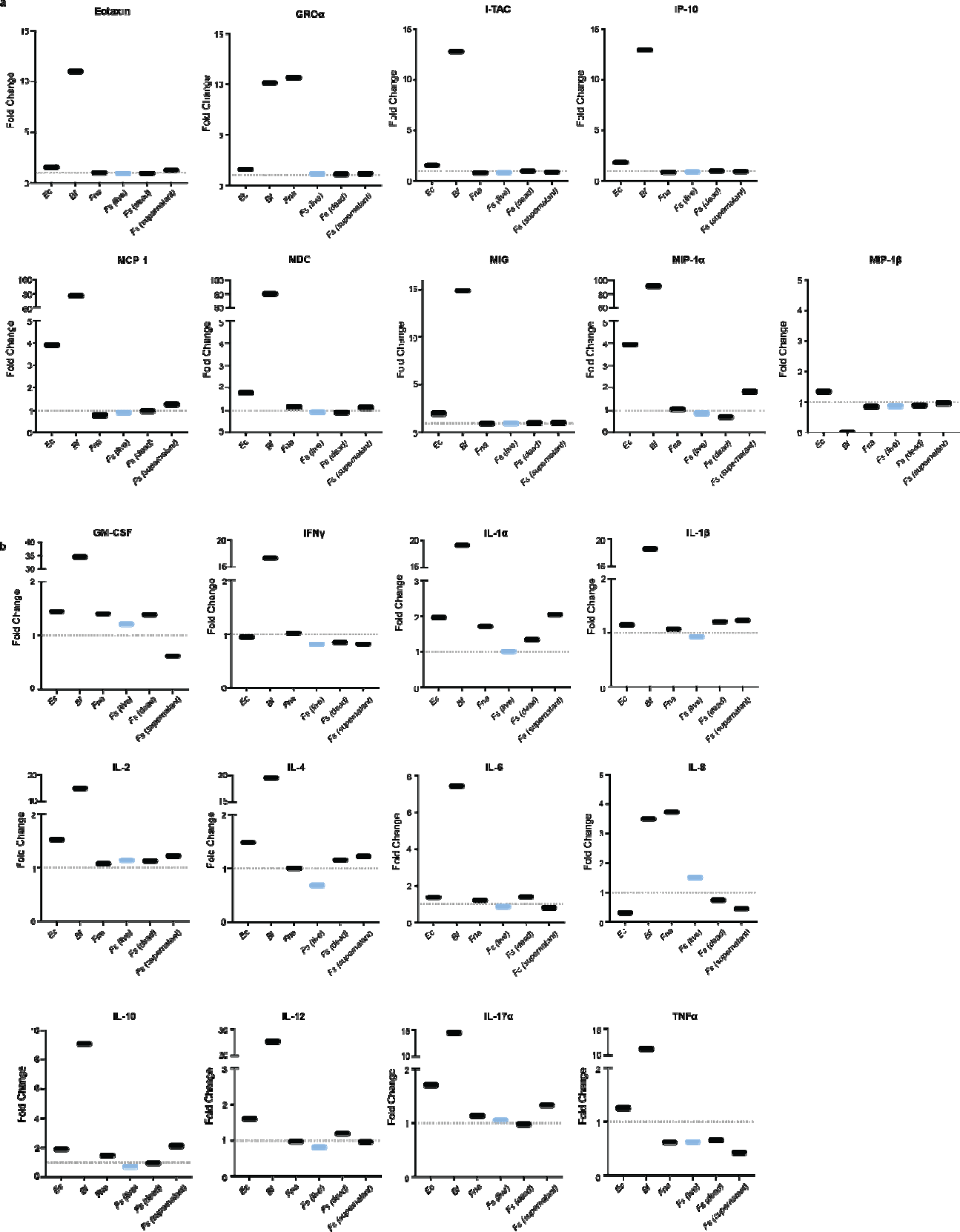
Cytokine and chemokine induction in HCT116 cells co-incubated with *F. sphaericum sp. nov*. Measurements of **a**) common chemokine and **b**) inflammatory cytokine levels measured via the Multi-Analyte ELISArray from HCT116 cells co-incubated with different *Fusobacterium* species at a multiplicity of infection (MOI) of 100 or antibody cocktail control, normalized to a no-bacteria control. Bacterial species used included *F. nucleatum* subsp. *animalis* (*Fna*) SB010 and *F. sphaericum sp. nov.* (*Fs*) SB021 live cells or oxygen-exposed dead cells. An additional co-incubation with *Fs* supernatant alone was also performed.

## Supplementary Table Legends

**Supplementary Table 1: RAST annotation of *F. sphaericum sp. nov.* SB021 genome**

The *F. sphaericum sp. nov.* SB021 genome was annotated by the <u>R</u>apid <u>A</u>nnotation using <u>S</u>ubsystem <u>T</u>echnology (RAST) pipeline. Table indicates resulting genes for each contig, with their length and predicted product.

**Supplementary Table 2: Methyl-modified motifs across *F. sphaericum sp. nov.* SB021 genome**

SB021 methyl-modified nucleotides were analyzed using Single-Molecule Real Time Sequencing (SMRTSeq)^10^ kinetics (Basemod analysis) and the predicted methyl-modified motif acquired via REBASE^39^ analysis.

**Supplementary Table 3: BLASTn analysis of *F. sphaericum sp. nov.* SB021 16S rRNA gene**

Table shows BLASTn top 100 results for the *F. sphaericum sp. nov.* SB021 16S rRNA gene by decreasing percent identity.

**Supplementary Table 4: Average nucleotide identity (ANI) scores between *Fusobacterium* genomes**

Table shows the pairwise average nucleotide identity (ANI) scores between each *Fusobacterium* genome (Table 1).

**Supplementary Table 5: GTDB-tk analysis of *F. sphaericum sp. nov.* SB021 genome**

Table shows the assigned phylogenetic classification of SB021 as determined by GTDB-tk^13^.

**Supplementary Table 6: Analysis of *F. sphaericum sp. nov.* SB021 anvi’o-identified gene clusters**

The *Fusobacterium* pangenome was characterized using the analysis and visualization platform for microbial ‘omics (Anvi’o)^19^. For each gene cluster (GC) identified in *F. sphaericum sp. nov*., the table shows its representative amino acid sequence, its mapped KO identifier by KEGG ortholog mapping via KofamKOALA^20^, and any identified hits by the virulence factor database (VFDB)^33^.

**Supplementary Table 7: CARD analysis of *F. sphaericum sp. nov.* SB021 genome**

Table shows results for the analysis of the *F. sphaericum sp. nov.* SB021 genome against the comprehensive antibiotic resistance database (CARD)^23^.

**Supplementary Table 8: SGB detection across human metagenomic samples**

Table shows relative abundance for *F. sphaericum sp. nov.* SB021 (SGB59307_group) and *Fusobacterium perfoetens* ATCC 29250 (SGB29302) across 21,382 human metagenomic samples. For each sample, dataset accession number and body site sampled is included.

## Methods

### Bacterial isolation and culturing

Written informed consent was obtained from patient as approved by the Fred Hutchinson Cancer Center Institutional Review Board. Primary adenocarcinoma tumor tissue was collected by hospital staff. Tissue was stored in sterile, chilled RPMI growth medium (Corning, Corning NY, USA) on ice. A small cross section of the tumor was cut from the main tissue mass, lightly scored, and then spread onto a small section on fastidious anaerobe agar (FAA) plates (Oxoid, ThermoFisher Scientific, Waltham, MA, USA) supplemented with 10% defibrinated horse blood (DHB; Lampire Biological Laboratories, Fisher Scientific, Pipersville, PA, USA). Using a sterile loop, the section was spread on the plates for the isolation of single colonies and subsequently, plates were incubated at 37°C under anaerobic conditions. Every 48 hours the plates were checked for new growth and the identity of colonies was determined by colony PCR. For all subsequent experiments, *F. sphaericum sp. nov* SB021 was grown for 48 hours on FAA+10% DHB plates supplemented with josamycin (3 μg/ml), vancomycin (4 μg/ml), and norfloxacin (1μg/ml) (Sigma Aldrich, USA) and incubated at 37°C under anaerobic conditions (AnaeroGen Gas Generating Systems, Oxoid, ThermoFisher Scientific, USA).

### 16S rRNA bacterial colony PCR

Single colonies were collected with a 1μl loop and suspended in 20μl of nuclease free water (Promega, Madison WI, USA). The same 1μl loop was then used to streak a FAA + 10% DHB plate to subculture and isolate pure colonies. The suspensions were placed in a thermocycler at 98°C for 20 minutes and then cooled to 4°C. 2μl of the lysate was subsequently used in a 20μL PCR reaction (Promega, Madison WI, USA) with forward primer 342F (5’- CTA CGG GGG GCA GCA G -3’) and reverse primer 1492R (5’- TAC GGY TAC CTT GTT ACG ACT T -3’) at a final concentration of 1μM. PCR products were shipped for Sanger Sequencing and sequencing results were analyzed using NCBI’s BLASTn for species level identification. Identified strains were stocked in 60% TSB with 40% glycerol (v/v) at −80°C.

### High-molecular weight genomic DNA (gDNA) extraction

High molecular weight genomic DNA was extracted using the MasterPure^TM^ Gram Positive DNA Purification Kit (Epicentre, Lucigen, USA). Cells from two plates were re-suspend in 1.5mL 1X PBS and harvested by centrifugation. Pellets were subjected to manufacturer instructions modified by doubling all reagent volumes and removing vortexing steps to prevent DNA shearing. HMW gDNA was quantified using a Qubit fluorometer (ThermoFisher Scientific, USA).

### PacBio Single-Molecule Real-Time Sequencing and Genome Assembly

Single molecule real-time sequencing (SMRT-Seq)^10^ was carried out on a PacBio Sequel instrument (Pacific Biosciences, USA) or a PacBio Sequel II instrument (Pacific Biosciencies, USA) at the University of Minnesota Genomics Center. Sequencing reads were processed using Microbial Assembly pipeline within Pacific Biosciences’ SMRTAnalysis pipeline version 9.0.0.92188. Additional assembly was performed using Flye assembler version 2.8 (https://github.com/fenderglass/Flye). Functional predictions for genes was conducted with the Rapid Annotations using Subsystems Technology (RAST)^40,41^ tool (**Supplementary Table 1**).

### Phylogenetic classification

The full-length 16S rRNA gene sequence for *F. sphaericum sp. nov.* SB021 was extracted from its whole genome sequence and analyzed using NCBI BLASTn (**Supplementary Table 3**). For available *Fusobacterium* type strains, the 16S rRNA gene sequence was downloaded and all resulting sequences were aligned via MEGA X^42^ using the MUSCLE clustering algorithm from which a maximum-likelihood dendrogram was generated. kSNP3^12^ with a kmer size of 17, resulting in a fraction of core kmers (FCK) of 0.113, was used to generate a maximum-likelihood whole-genome, reference-free phylogeny of these strains. Final tree were generated using the interactive tree of life (iTOL) tool, version 5^43^. Average nucleotide identity was calculated using the EZ BioCloud ANI calculator^44^. Further phylogenetic classification was determined using GTDB-tk^13^ (https://github.com/Ecogenomics/GTDBTk). For *Fusobacterium nucleatum*, ANI thresholds have led to adoption of different naming schemes based on recognition of the historical four subspecies as distinct species. In this work, we have implemented the *F. nucleatum sensu latu* nomenclature, referring to each subgroup by their historical subspecies name.

### Pangenomic Analysis

Pangenome analysis was performed using the <u>An</u>alysis and <u>Vi</u>sualization platform for microbial <u>‘o</u>mics (Anvi’o) workflow^19^ with standard thresholds set to a minbit of 0.5 and an MCL of 2. All *F. sphaericum* sp. *nov.* gene clusters were analyzed via KofamKOALA^20^ for KEGG ortholog mapping for their putative functions and analyzed against the Virulence Factor Database (VFDB)^33^ to identify putative virulence-associated genes.

### Bacterial Gram Staining

Gram staining was performed on *F. sphaericum sp. nov.* SB021 uniform bacterial smears after growth at 37°C for 48 hours followed by toxic oxygen exposure using a Gram Stain Kit (Remel, Lenexia, KS, USA). The stains were imaged with TissueFAXS microscope system (TissueGnostics, Vienna, Austria) with 100x oil immersion at Fred Hutchinson Cancer Center, Seattle, WA, USA.

### Biolog PM10 phenotype and anaerobe identification test panel microarray plates

*F. sphaericum sp. nov.* SB021 and *F. nucleatum* subsp*. animalis* SB010 were grown on FAA plates (Oxoid, Thermo Fisher Scientific) supplemented with 10% DHB (Fisher Scientific) incubated at 37°C in a Concept1000 anaerobic chamber (BakerRuskinn) for 24 h. PM10 and AN plates with corresponding inoculating fluids (IF-0a/IF-10b, and AN-IF, correspondingly) were pre-reduced, either at 4°C overnight (PM10) (AnaeroGen Gas Generating Systems, Oxoid, Thermo Fisher Scientific) or by manufacturer (AN). PM10 and AN plates were brought to room temperature under anaerobic conditions (AnaeroGen Gas Generating Systems, Oxoid, Thermo Fisher Scientific). Bacterial suspensions were made and added to each plate under anaerobic conditions at 37°C (BakerRuskinn). For PM10 plates, bacterial cells were resuspended in 2 ml of pre-reduced IF-0a and normalized across all samples to an optical density at 600nm (OD_600nm_) of 0.179 as recommended by Biolog. The final suspension was prepared by combining 0.75 ml of normalized bacterial suspension with 11.25 ml of mix B (100 ml pre-reduced IF-10b with 1.2 ml dye mix D, and 11.18 ml pre-reduced sterile water) to a final volume of 12 ml. For AN plates, bacterial cells were resuspended in 3 ml of pre-reduced AN-IF and normalized across all samples to an OD_600nm_ of 0.1 as recommended by Biolog. For both PM10 and AN plates, 100 μl of their respective final suspension was added to each well. All plates were then equilibrated to aerobic conditions at room temperature for 10 min and then incubated under anaerobic, hydrogen-free conditions for 24 h at 37°C (AnaeroGen Gas Generating Systems, Oxoid, Thermo Fisher Scientific). All plates were imaged and absorbance at 590 nm was quantified using a plate reader (Biotek).

### API 20A and API Zym Metabolic Profiling

The metabolic profile of *F. sphaericum sp. nov.* SB021 was determined with the API 20A Gallery System (bioMérieux, Marcy-l’Étoile, France) by utilization of colonies after 24h anaerobic culture. The colonies were resuspended in phosphate-buffered saline (PBS, Corning, Corning, NY, USA) to obtain an optical density at 600nm (OD_600nm_) of 0.4-0.5 (=3 McFarland) for inoculation of pre-coated cupules on the incubation strip. Results were read after 24 hours and additional procedures were carried out as indicated by the manufacturer. The enzymatic profile was investigated similarly with the API Zym Gallery System (bioMérieux, Marcy-l’Étoile, France), except for an OD_600nm_ of 0.7 (=5-6 McFarland) and an incubation period of 4-4.5 hours. Both API 20A and API Zym were performed with 3 replicates.

### Antibiotic Sensitivity Testing

*In vitro* testing of antibiotic susceptibility of *F. sphaericum sp. nov.* SB021 was performed on harvested bacteria after 24 hours incubation at 37°C under anaerobic conditions. Cells were resuspended in PBS to an OD_600nm_ of 0.5 and used to seed bacterial lawns on fastidious anaerobe agar (FAA) plates (Oxoid, ThermoFisher Scientific, Waltham, MA, USA) supplemented with 10% defibrinated horse blood (DHB; Lampire Biological Laboratories, Fisher Scientific, Pipersville, PA, USA). Antibiotic reading strips (MIC test strips; Liofilchem Diagnostics, Waltham, MA, USA) were placed on the lawns with sterile tweezers. All experiments were performed under anaerobic conditions and in triplicates for each antibiotic agent including negative controls (no bacteria). Results were read after 24 hours of incubation.

### Mammalian Cell Culture

Mammalian cell culture was performed on the cell lines HCT116 (CCL-247; ATCC, Manassas, VA, USA) and HT-29 (HTB-38; ATCC, Manassas, VA, USA). Mammalian cells were grown in T75 culture flasks using McCoy’s 5A culture medium (Iwakata and Grace Modification) (Corning, Corning NY, USA) with 10% fetal bovine serum (FBS; Sigma-Aldrich, St. Louis, MO, USA), and antibiotics (1x Pen/Strep: penicillin (5000 units/ml), streptomycin (5000 µg/ml), Gibco, Carlsbad, CA, USA) and incubated constantly in a 37°C/5% CO_2_ air humidified incubator. Cells were checked microscopically for monitoring purposes (growth rate, confluency) and split upon reaching confluency (>=80%). Trypsin-EDTA (Gibco, Carlsbad, CA, USA) was added to the flask to dissociate cells from flask surface, followed by neutralization of the dissociation agent with growth medium. Subsequently, the cell suspension was centrifuged for 3 minutes at 302 RCF at room temperature. The cell pellet was resuspended in sterile medium and seeded into new flasks. Passage numbers were recorded and did not exceed *n*=10. Prior to co-culture experiments, cells were split as described above and resuspended in antibiotic-free sterile growth medium, added to 6-well plates to a final concentration of 1.5×10^6^ cells per well and incubated for 24 hours before experimental testing.

### Scanning Transmission Electron Microscopy (S/TEM)

*F. sphaericum sp. nov.* SB021 was cultured as described above. Cell size of *F. sphaericum sp. nov.* SB021 was determined via scanning transmission electron microscopy (S/TEM). We co-incubated HCT116 human colorectal carcinoma cell line (CCL-247; ATCC, Manassas, VA, USA) at low passage (<10) with *F. sphaericum sp. nov.* SB021 (OD_600nm_ = 1.0) for 4 hours followed by fixation with ½ strength Karnovsky’s fixative (2.5% glutaraldehyde and 2% paraformaldehyde in 0.1M cacodylate buffer) and further sample preparation on the same day. Specimens were viewed on a JEOL JEM 1400 transmission electron microscope (JEOL, Tokyo, Japan) with an accelerating voltage of 120kV. Images were acquired with a Gatan Rio 4k x 4k CMOS Camera (Gatan, AMETEK, Berwyn, PA, USA) at Fred Hutchinson Cancer Center, Seattle, WA, USA.

### Confocal Microscopy of *Fusobacterium* co-cultures with human cell lines

*Fusobacterium* strains, *F. sphaericum sp. nov.* SB021 and *F. nucleatum* subsp. *animalis* SB010, were grown for 48 hours at 37°C in anaerobic conditions and resuspended in 1 mL of PBS at a final concentration of 5×10^8^ cells/mL. Bacterial membranes were stained in this suspension with 50 µl of 100 mg/mL FM 4-64FX (Molecular Probes) and washed twice in PBS by centrifugation for 7,000 x g at RT. Cells from human adenocarcinoma tumor cell lines, HCT116 and HT-29, were grown and passaged as described above and resuspended in 1 mL of McCoys 5A + 10% FBS. 6×10^4^ HCT116 or HT-29 cells were seeded with an MOI of 100 bacterial cells in 6-well plates. Bacteria were centrifuged onto HCT 116 and HT-29 at 300 x g for 5 minutes at room temperature (RT), and plates were immediately placed in a 37°C CO_2_ incubator and cultured together for 4 hours. Cell lines were then washed 3x with PBS (all washes are 5 min in duration at RT) and then fixed in 4% paraformaldehyde in PBS for 30 min at RT. Following fixation, wells were washed 3x in PBS and then permeabilized with 0.2% (*v/v*) Triton X-100 in PBS for 4 mins at RT. Cells were washed 3x in PBS and then stained for 20 min at RT with 2 drops/mL of NucBlue Fixed Cell Strain ReadyProbes (Invitrogen, Carlsbad, CA, USA) and ActinGreen 488 Ready Probes (Invitrogen) to stain DNA and actin, respectively. Cells were washed a final 3x with PBS and then the coverslips were mounted onto glass slides in ProLong Gold antifade mounting medium (Invitrogen). Samples were viewed with a Leica SP8 confocal laser-scanning microscope (Leica, Wetzlar, Germany) for image acquisition. Representative confocal micrographs of 1024 x 1024 pixels (pixel size: 103.3 nm) were acquired and assembled using Fiji (with Bio-Formats Plugin)^49^.

### Analysis of Chemokine and Cytokine Induction in HCT116

To determine the immunologic impact of *Fusobacterium* presence on HCT116 human colon epithelial cells, we utilized the Multi-Analyte ELISArray kits protocol version 1.4 from SABiosciences (Qiagen, Valencia, CA, USA) for common human chemokines and inflammatory human cytokines according to the manufacturer’s instructions. Tests were performed with the following samples after a 4-hour co-incubation with HCT116 (low passage, <10): *F. sphaericum sp. nov.* SB021 at MOI of 1:100 (live and oxygen-exposed), *F. sphaericum sp. nov.* SB021 cell-free supernatant, *F. nucleatum* subsp. *animalis* SB010 at MOI 1:100 (live), a negative control (HCT116 only) and a positive control (cocktail of antibodies). Supernatants were centrifuged at 1,000 x g for 10 minutes to remove any particulate material, filter sterilized and 50μl of each experimental sample were added to the array with the specific capture antibodies (IL-1α, IL-1β, IL-2, IL-4, IL-6, IL-8, IL-12, IL-17α, IFN-γ, TNF-α and GM CSF; IL-8, MCP-1, RANTES, MIP-1a, IP-10, I-TAC, MIG, Eotaxin, TARC, MDC, GROα), followed by an incubation at room temperature (RT) for 2 hours. Subsequently, the wells were buffer washed and 100μl of diluted biotinylated detection antibodies were added for a 1h-incubation at RT in the dark. After another washing procedure, the wells were filled with 100μl of Avidin-horseradish peroxidase (HRP) and incubated at RT for 30 minutes, again in the dark. We then added development and stop solutions to detect absorbance changes immediately after termination of the experiment with a microplate reader (Synergy H4 Hybrid Reader, BioTek) at 570nm and 450nm, visualized with Gen5 Synergy H4 software (Version 3.08, BioTek). Raw data of the absorbance reading were normalized to the cell counts of HCT116 cells incubated without bacteria. This experiment was executed in duplicates.

For quantification of immunologic effects, we utilized the Single-Analyte ELISArray kit protocol version 1.4. from SABiosciences (Qiagen, Valencia, CA, USA) for the chemokine IL-8 according to the manufacturer’s instructions. Co-cultures were set up as described above with *F. sphaericum sp. nov.* SB021 at MOI of 1:100 (live), *F. sphaericum sp. nov.* SB021 cell-free supernatant, *F. nucleatum* subsp*. animalis* SB010 at MOI 1:100 (live), *Escherichia coli* TOP10, or a positive control (HCT116 only) and incubated for 4-hours or 24-hours. Supernatants were centrifuged at 1000 x g for 10 minutes and filter sterilized subsequently. A serial dilution of the antigen standard was prepared and 50μl of each sample was added to the array, followed by an incubation for 2 hours at RT. Similar to the Multi-Analyte kit, the array was then incubated with detection antibodies and subsequently Avidin-HRP to determine the attachment of detection antibodies to the chemokine. After the color development, changes of absorbance were read at 570nm and 450nm and raw data were equally normalized. This experiment was executed in duplicates. Statistical comparison was calculated by applying a one-way ANOVA using GraphPad Prism v7.0 Software (GraphPad Software).

### Analysis within Human Metagenomic Specimens

Human metagenomic samples were profiled at SGB-level resolution with MetaPhlAn 4 (v4.beta.1) using the vJan21 markers database^45^ SGB assignment of *Fusobacterium sphaericum* sp. *nov.* SB021 (SGB6028, part of SGB59307_group) and *Fusobacterium perfoetens* ATCC 29250 (SGB29032) was performed with ‘phylophlan_metagenomic’ subroutine of PhyloPhlAn 3 (v3.0)^46^.

### Data Availability

*F. sphaericum sp. nov.* strain SB021 whole genome sequence is available in GenBank under the accession number SAMN15202580 and methylome is available in the Restriction Enzyme Database (REBASE) under organism number 39739.

## Acknowledgements

We would like to thank the Cellular Imaging Shared Resource of the Fred Hutchinson Cancer Center/University of Washington Cancer Consortium (P30 CA015704). Scientific Computing Infrastructure at Fred Hutchinson Cancer Center was funded by ORIP grant S10OD028685.

## Contributions

Y.E., H.H., C.D.J. and S.B. designed the study. M.Z.R, Y.E., F.E.D, C.D.J., and S.B. wrote the paper. A.B. and K.D.L. processed patient tissue specimen. A.B. and K.D.L. carried out microbial isolation and culture. M.Z.R., D.S.J. and C.D.J. performed DNA extraction, genome sequencing, and assembly. M.Z.R., Y.E., A.B., H.W., C.M., G.P., and S.S.M. performed computational and statistical analysis. Y.E., A.B., E.F.M., and K.D.L. carried out *in vivo* and *in vitro* assays. N.S., C.D.J., and S.B. obtained funding and supervised computational and wet lab experiments. All authors read and provided edits to the paper and contributed to the final version.

